# Multilayered regulation by RNA thermometers enables precise control of Cas9 expression

**DOI:** 10.1101/2025.05.12.651296

**Authors:** Elise K. Phillips, Dawn M. Klingeman, Adam M. Guss, Carrie A. Eckert, William G. Alexander

## Abstract

Cas9-based genome editing technologies can rapidly generate mutations to probe a diverse array of mutant genotypes. However, aberrant Cas9 nuclease translation and activity can occur despite the use of inducible promoters to control expression, leading to extensive cell death. This background killing caused by promoter leakiness severely limits the application of Cas9 for generating mutant libraries because of the potential for population skew. We demonstrate the utility of temperature sensitive RNA elements as a layer of post-transcriptional regulation to reduce the impact of promoter leak. We observe significant temperature-dependent increases in cell survival when certain RNA thermometers (RNATs) are placed upstream of the *cas9* coding sequence. We show that the most highly repressing RNAT, *hsp17rep*, significantly reduces population skew with a library of characterized guide RNAs (gRNAs). This information should be applicable to all Cas9-based methods and technologies.

**Notice:** This manuscript has been authored by UT-Battelle, LLC, under contract DE-AC05-00OR22725 with the US Department of Energy (DOE). The US government retains and the publisher, by accepting the article for publication, acknowledges that the US government retains a nonexclusive, paid-up, irrevocable, worldwide license to publish or reproduce the published form of this manuscript, or allow others to do so, for US government purposes. DOE will provide public access to these results of federally sponsored research in accordance with the DOE Public Access Plan (https://www.energy.gov/doe-public-access-plan).

**Graphical Abstract:** 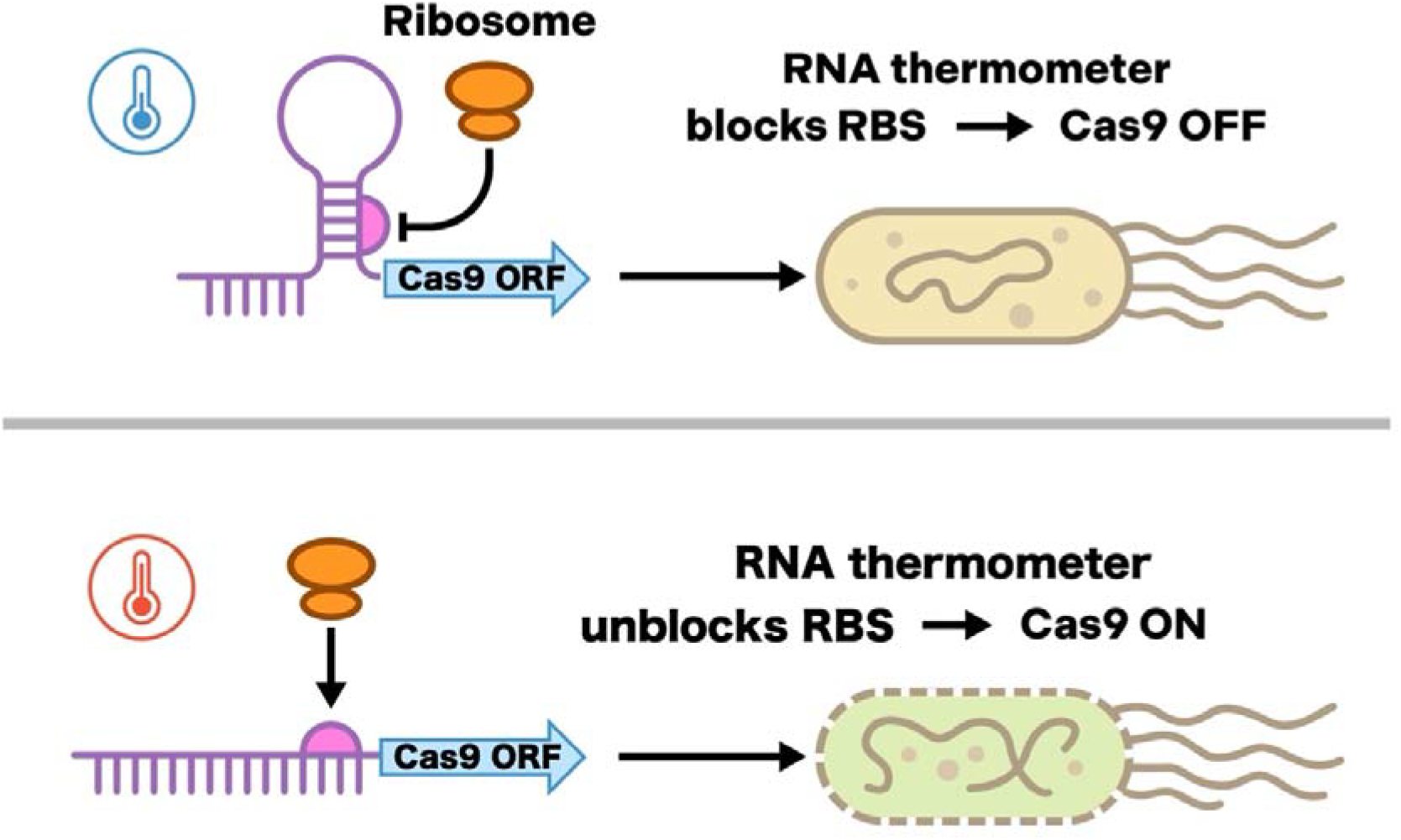

## Introduction

Controlled timing and degree of gene expression to optimally regulate desired traits is relevant to native hosts and genetically engineered organisms alike. One strategy for controlling gene expression is the use of inducible promoters, which alter gene expression in response to environmental stimuli such as carbon source or temperature. Commonly used bacterial inducible promoters, including the arabinose inducible promoter (P_araBAD_) (1), the lambda phage heat-inducible promoter (P_L_) (2), the lactose inducible promoter (P_lac_) (3), and the rhamnose inducible promoter (P_Rha_) (4), have different inducing conditions, dynamic ranges, and basal expression (5), which make them suited for different applications. However, inducible promoters often “leak,” meaning some level of transcription occurs in the absence of induction (6, 7). While the effects of this unintended expression is usually negligible, errant Cas9 expression is known to be problematic in heterologous contexts (8–12). For example, leaky Cas9 expression is thought to prevent gRNA library maintenance due to low levels of expressed Cas9 acting on library members, skewing the population stochastically. Additionally, the inherent intolerance of Cas9 expression in some microbes can preclude implementation in new organisms (13, 14). Therefore, identification of inducible promoters with tight regulation for Cas9 expression is crucial; unfortunately, even the most tightly regulated natural promoter known (the GAL1 promoter from *Saccharomyces cerevisiae*) exhibits detectable expression in repressive conditions (15–18). Natural gene expression networks have evolved multi-element control schemes to alter gene expression (19–22). For example, *Bacillus subtilis* regulates extracellular protease production via transcription factors that form feedback loops and whose activity is further modulated by the phosphorylation state of these transcription factors (23–26). Mimicking this naturally occurring layered control approach, synthetic biologists have developed genetic constructs that incorporate additional layers of post-transcriptional regulation, such as variable strength ribosome binding sites (27–29), riboswitches (12, 30), and small transcription activating RNAs (STARs) (31).

RNA thermometers (RNATs) are a group of RNA-based regulatory elements that inhibit translation in a temperature-dependent manner (32, 33). Simple RNATs are characterized by a stem and loop structure, commonly referred to as a “hairpin,” that sterically inhibits ribosome binding. At elevated temperatures base pairing in the stem becomes weaker causing the stem to melt open, derepressing protein expression. These regulatory elements are particularly appealing as their function is based solely on RNA structure and, thus, should be largely organism agnostic. Despite their obvious utility, application has been somewhat limited (34, 35), with no known use to date in the control of genome editing tools. Some barriers to their implementation include complex, challenging to predict structures, and open reading frame-inclusive RNATs (36–38). Thus, many RNATs can be challenging to engineer for generalization to novel systems, despite recent improvements in structure prediction models and software (39–41). Well characterized RNATs that function independently of the downstream protein sequence are therefore of particular interest. One such RNAT is present in the cyanobacteria *Synechocystis* sp. PCC 6803 upstream of the heat shock protein gene *hsp17* (42). Previously, this thermometer was mutated to alter its denaturation temperature and evaluated in both *Synechocystis* and *E. coli*. This previous characterization and proven tractability across species made *hsp17*-derived RNATs promising candidates to explore for modulating Cas9 expression.

In this work, we evaluated RNA thermometers for controlling Cas9 expression in *E. coli* by way of a simple *in vivo* survivorship assay (Figure 1) (see Methods)(9). Several tested RNATs significantly reduce Cas9 expression as demonstrated by improved survival in the presence of targeting gRNA vectors and a drastic reduction in the skew of library design abundances, enabling for Cas9 expression on demand. We demonstrated that a RNAT-containing, multilayered inducible expression system could be utilized to stably maintain pools of gRNAs, providing tight control of expression without significant loss of diversity during intervals consistent with routine plasmid preparations. These RNAT-based expression systems have the ability to provide a more accurate assessment of the starting population and more precise experimental control for pooled CRISPR-based genome editing strategies (43–49), as well as for CRISPR-based activation or interference (50–52).

**Figure 1:**
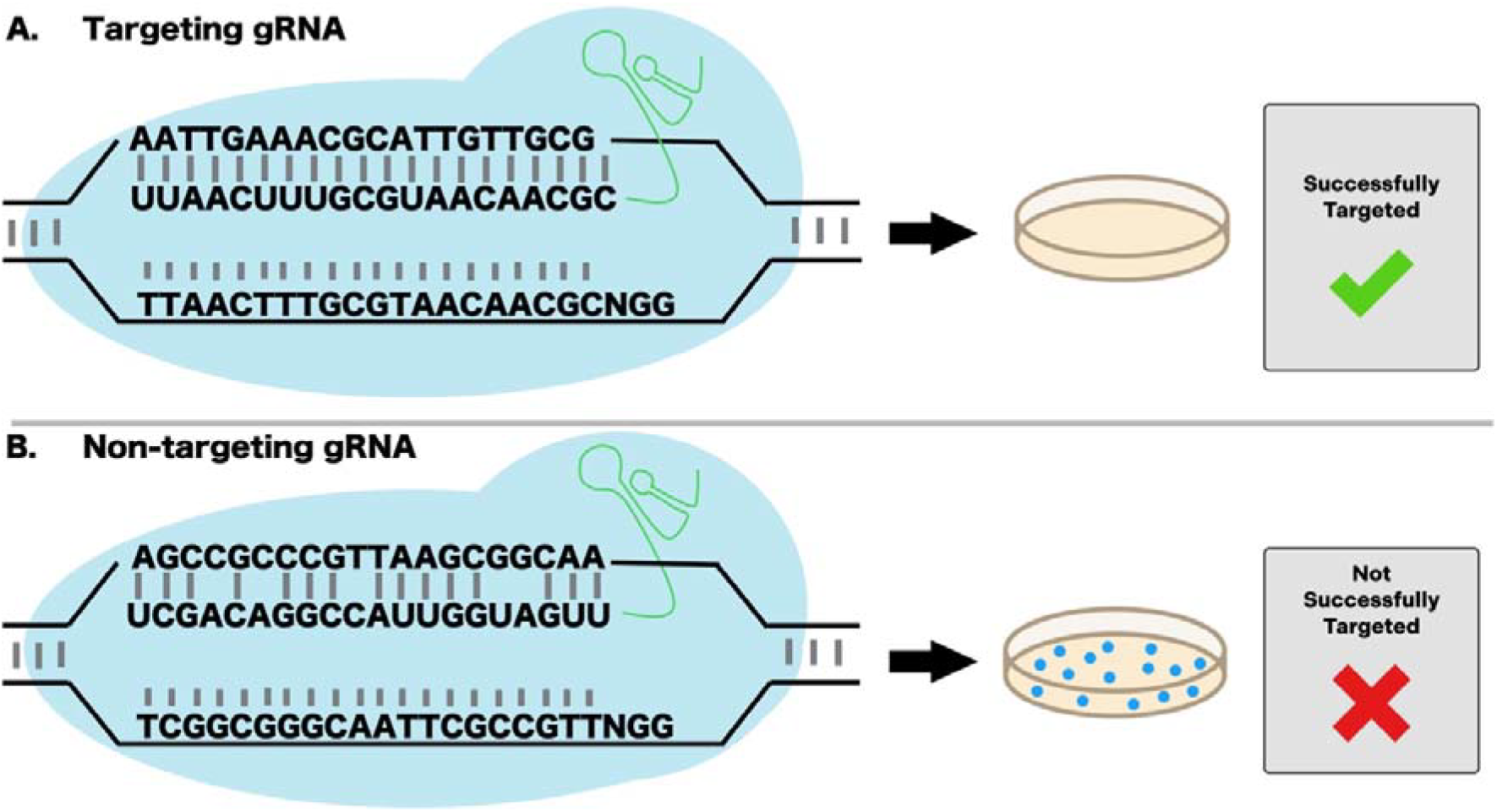
Strategy for calculating relative survivorship in Cas9-expressing cells containing targeting and non-targeting gRNAs. Relative survivorship is calculated by determining the ratio of colony forming units (CFUs) when cells are transformed with a chromosomally targeting **(A.)** or non-targeting **(B.)** gRNA. The targeting gRNA will result in a double strand break and death in the presence of Cas9. Conversely, the NT-gRNA has no recognition sequence targeting the genome and thus cells should survive with and without Cas9 induction. Relative survivorship ratios near one suggest low levels of Cas9 expression.

**Figure 2.**
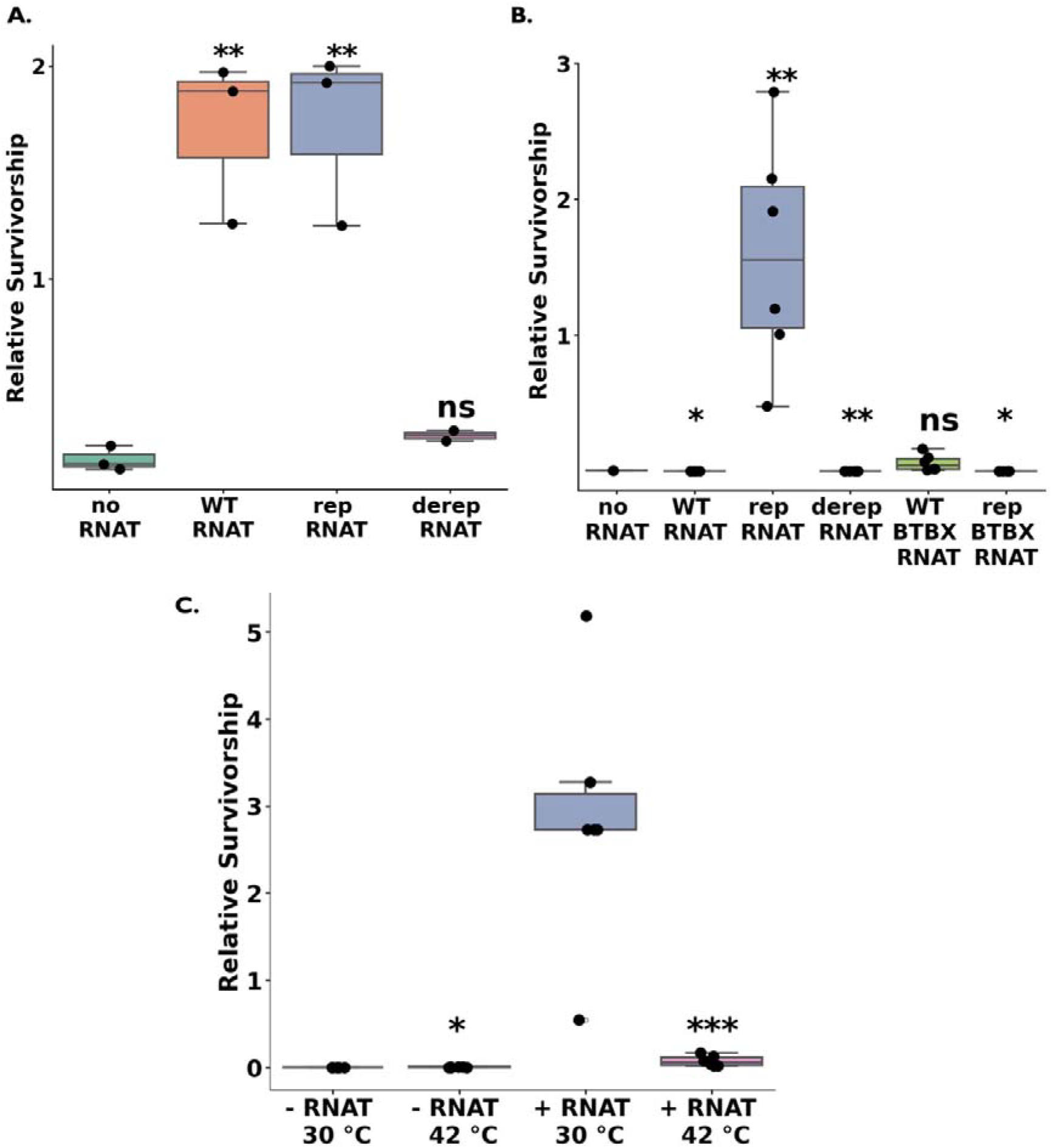
In combination with inducible promoters, RNA thermometers provide additive control of *cas9* expression. **A & B)** Relative survivorship for different promoter and RNA thermometer combinations with a targeting gRNA as determined by calculating fold change in CFUs of targeting gRNA compared to NT-gRNA with *cas9* expression driven by induction of the **A)** P_araBAD_ promoter, or **B)** P_L_ temperature sensitive promoter. **C)** Relative survivorship of cells grown with uninduced (30°C) or induced (42 °C) *cas9* with the P_L_ promoter with or without *hsp17rep* RNAT. Data represents 3 technical replicates of 1-2 biologic replicates. Significance determined by individual t-tests, where * 0.05 ≥ p > 0.01, ** 0.01 ≥ p >0.001, *** p ≤ 0.001. A) and B) make comparisons to no RNAT relative survivorship. C) Comparisons are made between 30 °C and 42 °C for each construct. Panels were generated in matplotlib (62) with packages pyplot, pandas v2.2.1 (63), numpy v1.26.4 (64), and seaborn v0.13.2 (65). Statistics were calculated using scipy v1.13.1 (66). Open circles denote data points lying outside of 1.5x the third quartile.

## Materials and Methods

### Bacterial Culture

Cells were cultured in LB-Lennox with antibiotics added as appropriate at the following concentrations: carbenicillin 100 µg/mL, chloramphenicol 15 µg/mL. Cells were grown with shaking at 250 rpm in 250 mL baffled flasks for experimental conditions and transformations, and 12 mL round bottom tubes for overnight cultures unless otherwise noted. Cells were grown at 30 °C unless they were shifted to 42 °C to induce *cas9* expression.

### Transformation of *E*. *coli* cells

Cells were grown at 30 °C to mid-log phase, washed three times with 10% cold glycerol (53) and electroporated with a MicroPulser (Bio Rad, Hercules, CA) using the bacteria Ec1 setting(1.8kV). After transformation, cells were recovered in super optimal broth (SOB) (54) for 1 hour at 30 °C.

### Vector construction

Inserts with approximately 18 base pair overlaps were cloned into vector backbones using NEBuilder HIFI DNA assembly master mix according to the manufacturer’s protocol (NEB Ipswich, MA). Gibson assembled plasmids were transformed into electrocompetent Lucigen E. cloni cells. Plasmids extracted using Monarch Plasmid Miniprep kit (NEB, Ipswich, MA) from four to eight clones were sequenced using Oxford Nanopore (Oxford, UK) sequencing R10.4.1 chemistry and sequence perfect clones were stored and used for subsequent experiments.

Sequences were assembled using Trycycler v0.5.5 (55), with initial assemblers Flye v2.9.4-b1799 (56) and miniasm v0.3-r179 (57). Confirmed plasmids were subcloned into electrocompetent *E. coli* MG1655 for survival and induction assays. Plasmids used in this study are listed in Supplemental Table 1. *hsp17* variant inserts were synthesized by Eurofins (Lancaster, PA) and pooled library inserts were synthesized by Genscript (Piscataway, NJ). All oligos used in this study are listed in Supplemental Table 2.

### Survival Assays: Uninduced Cas9

*E. coli* MG1655 cells containing *hsp17*-Cas9 vectors (pWGA121-123, pWGA182-184, pEKP13-14) were transformed with a plasmid containing either a non-targeting (pWGA128) or targeting (pWGA130) gRNA. Cells were recovered for one hour at 30 °C, serially diluted, then plated on LB carbenicillin + chloramphenicol and incubated overnight at approximately 28-30 °C. Cells possessing the targeting gRNA and Cas9 receive DSBs and die, while those with NT-gRNA survive. By dividing targeting gRNA CFUs by NT-gRNA CFUs, we determine relative survivorship. Relative survivorship values near one indicate Cas9 activity equal to the NT-gRNA, indicating absence of Cas9 expression (Figure 1).

### Survival Assays: Induced Cas9

Relative survivorship experiments with arabinose induction, pX2-Cas9 and derivative plasmids, were performed as described previously (58) with a few modifications. Briefly, cells were grown with 0.4% arabinose and at 30 °C rather than 37 °C.

For heat induction, cells were grown as described for the survival assays with the following modification: after 1 hour of recovery in SOB, the cells were incubated at 42 °C overnight, then plated on selective media to assess survival.

### Measuring gRNA library skew

A library of 500 functional and 500 non-functional verified gRNAs (59)was synthesized by Genscript (Piscataway, NJ). These libraries were cloned into pSS9 (44), then the gRNA plasmid library was extracted to produce the donor pool. Library coverage is reported in Supplemental Table 3. To gauge gRNA cutting activity, gRNAs with known non-targeting (pWGA128) and targeting (pWGA130) activity were added to the donor pool before transformation as “spike-in controls”. Equal masses of the donor plasmids pool were combined, using 100 pg of each control per 100 ng of pooled library plasmids. The plasmid libraries were then transformed into WA00135 (P_L_-Cas9) and WA00146 (P_L_-hsp17rep-Cas9). To calculate transformation efficiency, pWGA128 was individually transformed into both host strains, recovered for 1 hour at 30 °C, then plated. Additionally, the gRNA library transformations were diluted 50-fold in LB carbenicillin + chloramphenicol and incubated overnight (about 18 hours). 1.5 mL of the culture was pelleted, washed twice in an equal volume of DNase buffer (10 mM Tris, 2.5 µM MgCl_2_, 0.5 mM CaCl_2_, pH 7.5), then resuspended in 50 µL of buffer incubated with 5 µL of DNase1 at 37 °C for one hour to remove residual extracellular DNA. After DNase treatment, plasmids were extracted and frozen. A portion of remaining culture was diluted 50-fold to incubate an additional 24 hours (denoted day 2 in Figures 3 and 4). Plasmids from this passage were extracted and frozen as above. For the heat-activated library experiments, cells were grown at 30 °C for the 1^st^ day and transferred to 42 °C for the second.

**Figure 3:**
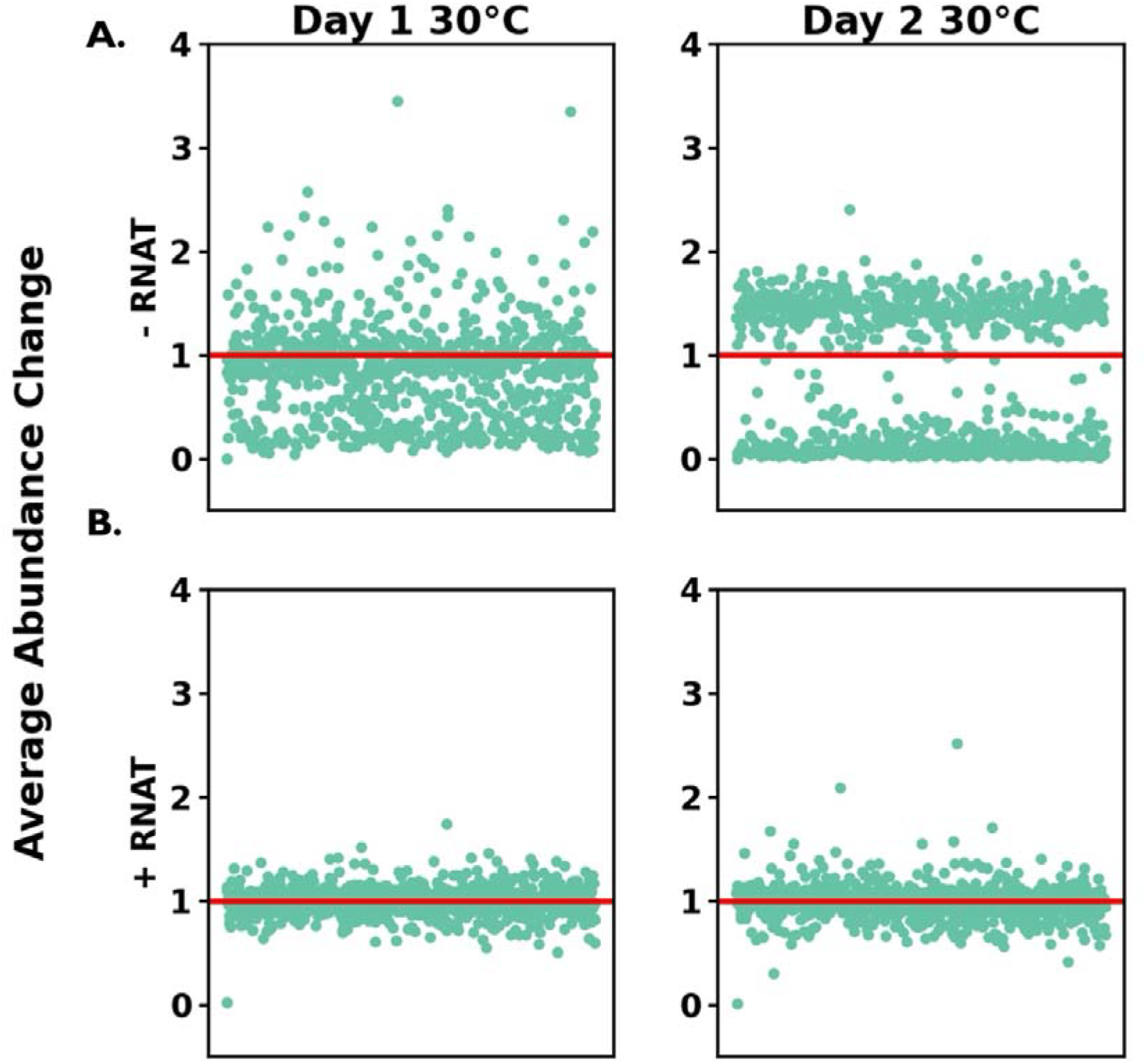
RNA thermometers prevent gRNA library skew by controlling *cas9* expression. Average change in abundance of gRNAs with uninduced *cas9* (30 °C) in the host compared to abundance in the donor pool. **A**. Without an RNAT. **B**. With the *RNAT hsp17rep*. gRNAs are evenly distributed across the x axis in alphabetical order according to their index name. Average abundance of three sequencing and two biologic replicates are shown. gRNAs with less than ten counts in the donor pool were excluded. Panels were generated in matplotlib (62) with packages pyplot, pandas v2.2.1 (63), numpy v1.26.4 (64), and seaborn v0.13.2 (65). Statistical significance was determined using student’s t-tests.

For library sequencing, the gRNA region of the plasmid was amplified using the phased primer approach as previously described using primers oWGA333-342(58). The resulting PCR products were subsequently gel purified. Amplicon DNA was prepared for Illumina sequencing using NEBNext Ultra II DNA Library Prep Kit for Illumina and sequenced with 3 technical replicates. Libraries were sequenced on an Illumina MiSeq (San Diego, CA) with V3 chemistry. A 600-cycle kit was used for paired end sequencing (2 × 151 × 8 × 8). PhiX control DNA was added to the run to increase based diversity in the amplicon pool. Paired-end reads were merged using the USEARCH -fastq_mergepairs function (60). gRNAs were then identified with USEARCH and counted using a custom python script. gRNAs with less than 10 counts in the donor pool were excluded from downstream analysis to prevent skew inflation due to their low abundance. gRNA abundance of the host populations was determined by dividing the host gRNA abundance by the average abundance of the corresponding gRNA in the donor pools (44). Student’s t-test was used to determine significant changes in gRNA abundance from the donor pool to the host.

## Results and Discussion

### Testing RNATs for multilayer control of Cas9 expression

To reduce leaky Cas9 expression, we were inspired by natural regulatory systems to install a second layer of Cas9 regulation by combining RNATs with frequently used inducible promoters (1–4). We compared the heat inducible promoter, P_L_, and the arabinose inducible promoter, P_araBAD_, with the *Synechocystis hsp17* RNATs described previously (42) as well as two ribosomal binding site (RBS) variants that we designed for this work. We observed significantly increased relative survivorship (Figure 1A) using the P_araBAD_ promoter in combination with both *hsp17WT* and *hsp17rep* RNATs (Figure 2A). However, with the P_L_ promoter, only the *hsp17rep* RNAT improved relative survivorship significantly (Figure 2B). The *hsp17derep* RNAT, which is less repressive than *hsp17WT*, did not improve survival with either promoter at 30 °C, which is consistent with previously reported results (42). Because exchanging RBSs can be used to fine-tune protein expression we probed the impact of altering the RBS on *cas9* expression by mutating the native *Synechocystis* RBS to the commonly used *E. coli* RBS *BTBX-1* (61) for a few of the P_L_ regulated constructs. When the stem sequence of *hsp17WT* and *hsp17rep* was mutated to include *BTBX-1* and its complement we observed a minimal increase in survival with the RNAT analogous to the *hsp17WT* RNAT but not the one analogous to the *hsp17rep* RNAT (Figure 2B). However, the *hsp17WT: BTBX-1* did not increase survival to the same magnitude as the initial *hsp17* RNAT variants.

With *P*_*L*_*-hsp17rep* identified as the most effective combination available for control of *cas9* expression, we tested its ability to control *cas9* expression in a temperature-dependent manner by examining relative survivorship. We observed significantly reduced survival at 42 °C with the targeting gRNA plasmid with the RNAT (Figure 2C), confirming the functionality of this *cas9* expression construct. Without the RNAT, we do not observe reduced survival at 42 °C, but we predict that this is due to the widespread cell killing from leaky *cas9* expression (Figure 2C). This temperature-dependent survival demonstrates the utility of a these RNATs as a secondary level of regulation for Cas9 and other potentially toxic genes.

### Utilizing RNATs for control of *cas9* expression for gRNA pooled libraries

Previous research utilizing gRNA libraries has operated under the assumption that leaky *cas9* expression can alter the relative abundances of a pool of gRNAs before “time zero” of the intended experiment(59). Thus, long-term storage of these transformed libraries in freezers has been avoided in favor of immediately placing constructed libraries under selection after transformation. While this provides reduced skewing of gRNA abundances during outgrowth from a freezer, these procedures are labor intensive, hamper reproducibility, and reduce the number of conditions a given gRNA pool can be tested against due to synthesis and experimental constraints.

Stable host storage circumvents the need to transform the pool *de novo* for each use, allowing for these transformed gRNA pooled libraries to be propagated from freezer stocks, providing more accurate measurements of the starting population prior to induction at the start of an experiment.

To evaluate whether RNA thermometers can be utilized to prevent gRNA library skew and control induction for a pooled gRNA library, we designed and synthesized a library containing one thousand unique gRNAs with previously validated functionality(59), divided evenly between known functional and non-functional gRNAs. This library was transformed into host *E. coli* strains where *cas9* was regulated by the P_L_ promoter in tandem with or without the RNAT *hsp17rep*.

These strains were grown under *cas9* repressive conditions (30 °C) and evaluated over 2 days (Figure 3). Without the RNAT, we observe more gRNAs with statistically significant changes in abundance on day 1, and an even greater number on day 2 (Figure 3A, Supplemental Data 1). This indicates population skew resulting from leaky *cas9* expression in the presence of the gRNA library. The two distinct subpopulations above and below an average abundance change of one (Figures 3 & 4, red line) correspond to non-targeting and targeting gRNAs, respectively, demonstrating the depletion of targeting gRNAs from the population in the absence of the RNAT. Conversely, the population with *cas9* additionally controlled by the RNAT showed a tighter distribution of gRNAs throughout day 1 and day 2, as well as a clustering around an average abundance change of one (Figure 3B), indicative of little to no change from the starting population of gRNAs. Further, without the RNAT the degree of the change from the donor population is more substantial on day 1 than with the RNAT. Without the RNAT the average change in abundance for each gRNA is approximately 40%, while there is only 13% change in abundance with the RNAT present. This becomes more pronounced on the second day where the average abundance change is 64% and 11% without and with the RNAT respectively. This indicates that the *hsp17rep* RNAT in combination with an inducible promoter can reduce gRNA population skew to minimal levels by allowing for tighter control of *cas9* expression.

We then wanted to confirm that heat-induction of the RNAT controlled *cas9* expression cassette would lead to library skew due to active Cas9 in the presence of the gRNA library. We again performed a two-day incubation of cells transformed with gRNA libraries, however, in this case, the first day was incubated at 30 °C (non-inducing), and the cells were shifted on day 2 to 42 °C (inducing). We observed similar population trends without the RNAT at 30 °C with skew increasing over time and at 42 °C (Figure 4A, Supplemental Data 2). In the presence of the RNAT, we saw results consistent those in Figure 3B for day 1 at 30 °C (Figure 4B). However, when shifted to inducing conditions at 42 °C in day 2, we observe skew develop in the library (Figure 4B). The observed skew at inducing temperatures with the RNAT is similar to the amount of skew present on day 1 for -RNAT cells at 30 °C. This indicates activation of *cas9* expression and Cas9 cutting of the chromosome. The amount of library skew that occurs during active Cas9 expression with the RNAT at 42 °C is less pronounced than without the RNAT, which is likely because the population without the RNAT had already begun to skew on day 1. Together, these data show that *hsp17rep* is effective at preventing gRNA library skew at nonpermissive temperatures and enables gRNA library abundance changes at permissive temperatures.

**Figure 4:**
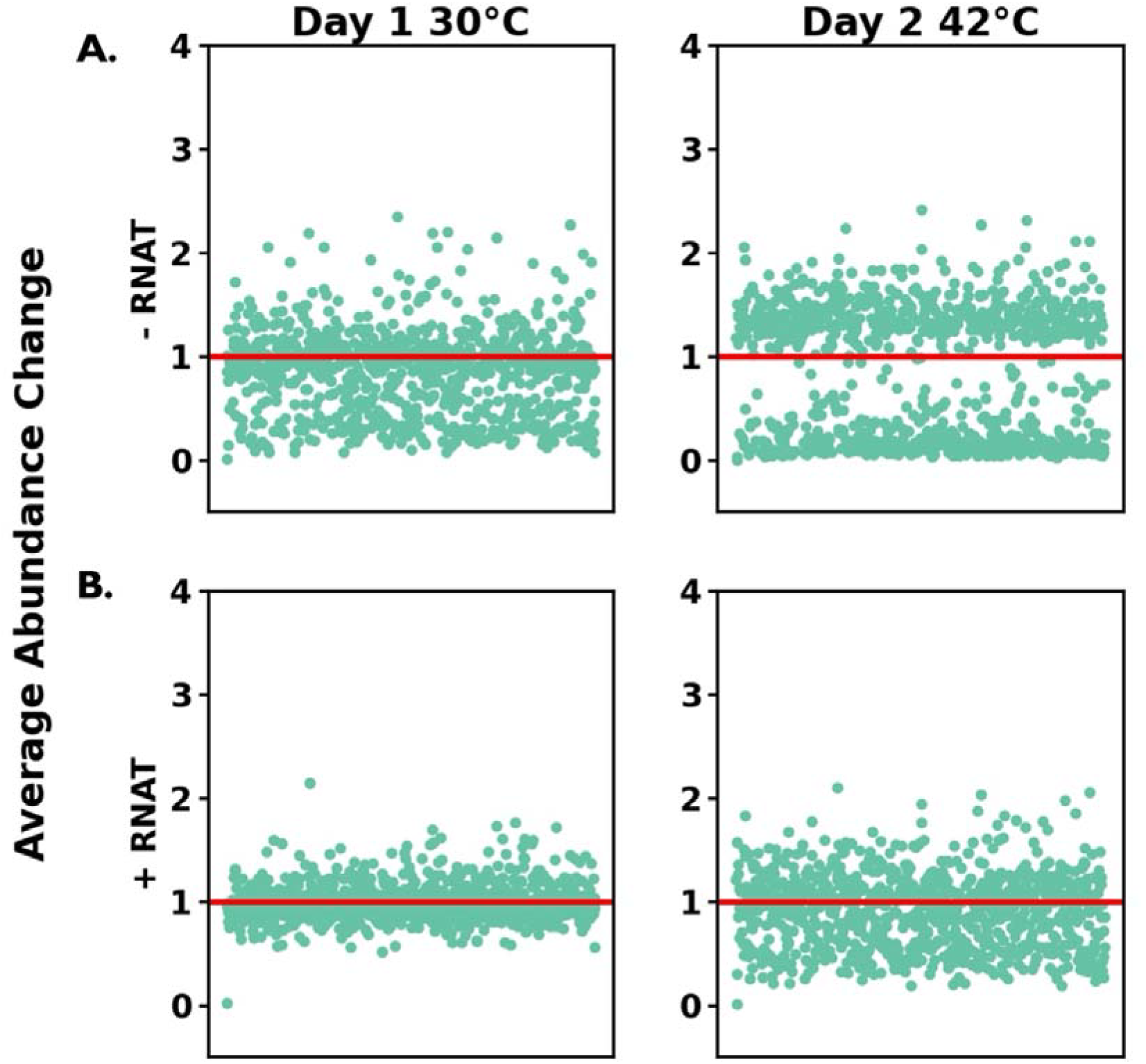
RNAT temperature induction of *cas9* expression leads to gRNA skew due to Cas9 cutting of the genome. Average change in abundance of gRNAs with uninduced (30 °C) and induced (42 °C) *cas9* in the host compared to abundance in the donor pool. **A**. Without an RNAT. **B**. With the *RNAT hsp17rep*. gRNAs are evenly distributed across the x axis in alphabetical order according to their index name. Average abundance of three sequencing and two biologic replicates are shown. Guides with less than ten counts in the donor pool were excluded. Panels were generated in matplotlib (62) with packages pyplot, pandas v2.2.1 (63), numpy v1.26.4 (64), and seaborn v0.13.2 (65). Statistical significance was determined using student’s t-tests.

## Conclusion

Driving Cas9 expression with leaky inducible promoters, while necessary, has required technical workarounds that hamper Design Build Test Learn (DBTL) cycles and potential reproducibility. We show that RNA thermometers derived from the *Synechocystis hsp17* RNAT are an effective element for reducing promoter leak and preventing premature *cas9* expression in *E. coli*. The tighter control of Cas9 activity prevents unwanted gRNA depletions in diverse libraries. *hsp17rep* prevents gRNA library skew over two days, making it useful for any large gRNA library, whether for editing, CRISPRi, or CRISPRa. If a specific application requires higher levels of protein expression than afforded by our construct, further optimization may be necessary. This RNAT-based approach could potentially be implemented for eukaryotic systems, as the functionality of the RNAT is based solely on the RNA structure thus making it highly translatable to other systems. Additionally, RNATs could be used to titrate expression of proteins of interest, such as those used in recombineering or proteins that are toxic to cells even in low concentrations.

## Contributions, Funding and Additional Information Author Contributions

Elise K. Phillips: Methodology, Writing-Original draft. Dawn M. Klingeman: Methodology, Writing-Reviewing and Editing. Carrie A. Eckert: Writing-Reviewing and Editing. William G. Alexander: Conceptualization, Methodology, Writing-Reviewing and Editing

## Acknowledgements

This research used resources of the Compute and Data Environment for Science (CADES) at the Oak Ridge National Laboratory, which is supported by the Office of Science of the U.S. Department of Energy under Contract No. DE-AC05-00OR22725. We would also like to acknowledge ORNL Creative Services for assistance in figure development.

## Supplemental Information

Supplementary Data and Tables are available at NAR online. Associated sequencing data can be found under NCBI’s BioProject PRJNA1260935. Plasmid GenBank accession numbers can be found in Supplemental Table 1.

